# Principles for engineering microbial composition and ecosystem functioning

**DOI:** 10.1101/730556

**Authors:** Michaeline B.N. Albright, Sanna Sevanto, La Verne Gallegos-Graves, John Dunbar

## Abstract

Microbial probiotics are designed to improve functions in diverse ecosystems, yet probiotics often fail to have the desired beneficial effects. The introduction of probiotics to an environment with a preexisting microbiome is analogous to an invasion event, but is rarely considered in this light. Here, we tested the relative importance of propagule pressure (inoculation dose and frequency) compared to biotic interactions (composition of introduced and resident communities) in driving microbial composition and functional outcomes following microbial community invasions in experimental microcosms. Ecosystem functioning was assessed through measurements of CO_2_ production and DOC (dissolved organic carbon) accumulation, an activity and an environmental modification metric, respectively. Further, to test the dependence of propagule pressures versus biotic interactions was dependent on environmental context, experiments were performed on two different substrates, R2A agar and plant litter. In both environments, we found that biotic interactions were more important than propagule pressure in driving microbial composition. Moreover, bacteria were more successful invaders than fungi. While successful invasion is a first step, ultimately the success of microbial invasions in microbiome engineering applications is measured by the impact on ecosystem functioning. As with shaping the microbiome composition, biotic interactions were key to functional outcomes, but the magnitude of the functional impact varied by environment. Identifying general principles that determine the community composition and functioning following microbial invasions is key to efficient community engineering.

**Significance:** With increasing frequency humans are introducing new microbes into pre-existing microbiomes to alter functioning. Examples include, modification of microflora in human guts for better health, and soil for food security and/or climate management. Probiotic applications are often approached as trial-and-error endeavors and have mixed outcomes. We propose that increased success in microbiome engineering may be achieved by better understanding of microbial invasions. We conducted a microbial community invasion experiment, to test the relative importance of propagule pressure and biotic interactions in driving microbial community composition and ecosystem functioning in microcosms. We found that biotic interactions were more important than propagule pressure in determining the impact of microbial invasions. Furthermore, the principles for community engineering vary among organismal groups (bacteria versus fungi).

## Introduction

With increasing frequency, humans are manipulating microbial communities by introducing microbes to new locations with preexisting microbial communities to achieve desired functional outcomes (1). Examples include modification of microflora in human guts for better human health, bioreactors for fuel production, and soil for improved plant performance and/or climate management (1–9). Feasibility of this approach depends on the predictable establishment and persistence of inoculated microbial communities. However, the parameters for successful introduction of communities are not well defined. Some microbiome engineering efforts have been successful, but often results fall short. Introduced communities may not establish or they may establish temporarily (1, 10). For example, the measured effects of probiotics are often short-lived in studies of mouse gut microbiota, suggesting that the introduced bacterial members are transient (11, 12). If establishment is successful the invaders may produce the desired impact on functioning, but alternatively they may have no impact, or the impact may even be negative. Overall, microbial community manipulations continue to be a trial-and-error endeavor with low success rates, pointing to the need to identify fundamental principles for successful microbiome engineering.

Microbiome engineering is related to plant and animal invasion biology, which generally aims to counteract invasive plant and animal species whereas microbiome engineering aims to promote invasions. In invasion biology, invasion success is determined by three main processes: 1) dispersal filtering; 2) environmental filtering; and 3) biotic filtering (13, 14). Dispersal determines which species spread to novel areas, either through natural or human mediated movement. Propagule pressure, a measure of dispersal used to describe the magnitude and pattern of the arrival of invasive individuals (15), is one of the most commonly tested factors shaping invasion success (10, 15–22). Environmental filtering occurs when the conditions of the new habitat filters species based on ecological niches and physiological adaptations. Invasion success increases if species are preadapted to an environment (23). Lastly, biotic interactions determine if introduced species fail to establish (biotic filtering) or successfully coexist with residents either by exploiting unused resources or by replacing residents through competition. Research in this arena has found that characteristics of resident communities, such as richness and diversity can enhance community resilience to invasion (1, 11, 24–27).

The relative importance of the three processes is unclear (14). Furthermore, success is measured as the establishment of an invasive species, but the impact of invaders on the composition of the larger community and the functioning of the ecosystem they invade is often not assessed (28, 29). And, in contrast to macroecology, studies examining factors influencing microbial invasion are still relatively rare (20, 30). Understanding the relative importance of dispersal, environmental and biotic filtering in the establishment of microbial invaders is likely to improve outcomes in diverse microbiome engineering applications. This is crucial as markets for human and animal probiotic products are expected to exceed $50 billion by 2022 (31) and increasingly plant probiotics are also being developed (32–34).

A promising approach to quantify factors leading to successful invasions is to introduce whole communities rather than individual taxa (35–38). Introducing complex communities provides the opportunity to perform many invasion tests simultaneously, instead of the more commonly applied sequential tests with macro and micro-organisms that typically introduce only one or a few invasive species at a time (15, 21, 39, 40). Furthermore, this approach recapitulates some types of microbial dispersal events, which can involve deposition of many disparate taxa into a novel location at same time, for example through rain (41). The concept of interactions or interchanges of entire microbial communities at ecosystem transition zones (42–44), coined microbial community coalescence, has been explored in a number of arenas (5). Here factors such as environmental conditions, mixing ratios, and temporal dynamics are hypothesized to influence microbial composition (5). Recent research also suggests that macro-organism invasions often involve multiple invasive species (45). As multi-species manipulations are often unfeasible with macro-organisms due to the large scale, this is a major arena for microbial ecology to contribute to invasion theory, just as rapid bacterial growth rates were advantageous for discoveries in evolutionary biology (46).

Here, we conducted a microbial community invasion experiment to test the relative importance of propagule pressure (delivery parameters) and biotic interactions on microbial community composition and ecosystem functioning. We manipulated four factors: 1) inoculation dose; 2) inoculation frequency; 3) invader microbial community and; 4) resident microbial community in a microcosm experiment. Since our goal was to decipher common rules for disparate microbiome engineering applications, manipulations were performed in microcosms with two distinct substrates. One environment was R2A agar and the other was plant (*Pinus Ponderosa*) litter on sand. R2A agar contains more labile carbon (C) and allows for more homogenous mixing, while the litter environment contains more recalcitrant C and greater structural complexity. We preadapted invaders to each environment, eliminating environmental filtering as a test factor. During *Phase I* of the experiment, we established diverse model microbial communities in microcosms by inoculating replicate microcosms with soil suspensions (Figure 1a). In *Phase II*, we conducted experimental manipulations adding invader microbial communities (also generated in *Phase I*) at different doses and frequencies to established resident microbial communities (Figure 1b). During *Phase II* we measured carbon dioxide (CO_2_) production and dissolved organic carbon (DOC) accumulation as our metrics of ecosystem functioning. While desired functional outcomes may vary widely depending on the application of interest, we chose CO_2_ production as a metric of microbial activity and DOC accumulation as a metric of microbial modification of the environment. We hypothesized that propagule pressure would play a larger role than biotic interactions in shaping microbial composition and ecosystem functioning. Propagule pressure is widely considered a primary determinant of invasion success (14, 47) and thus likely also has impacts on community composition and ecosystem functioning.

**Figure 1.**
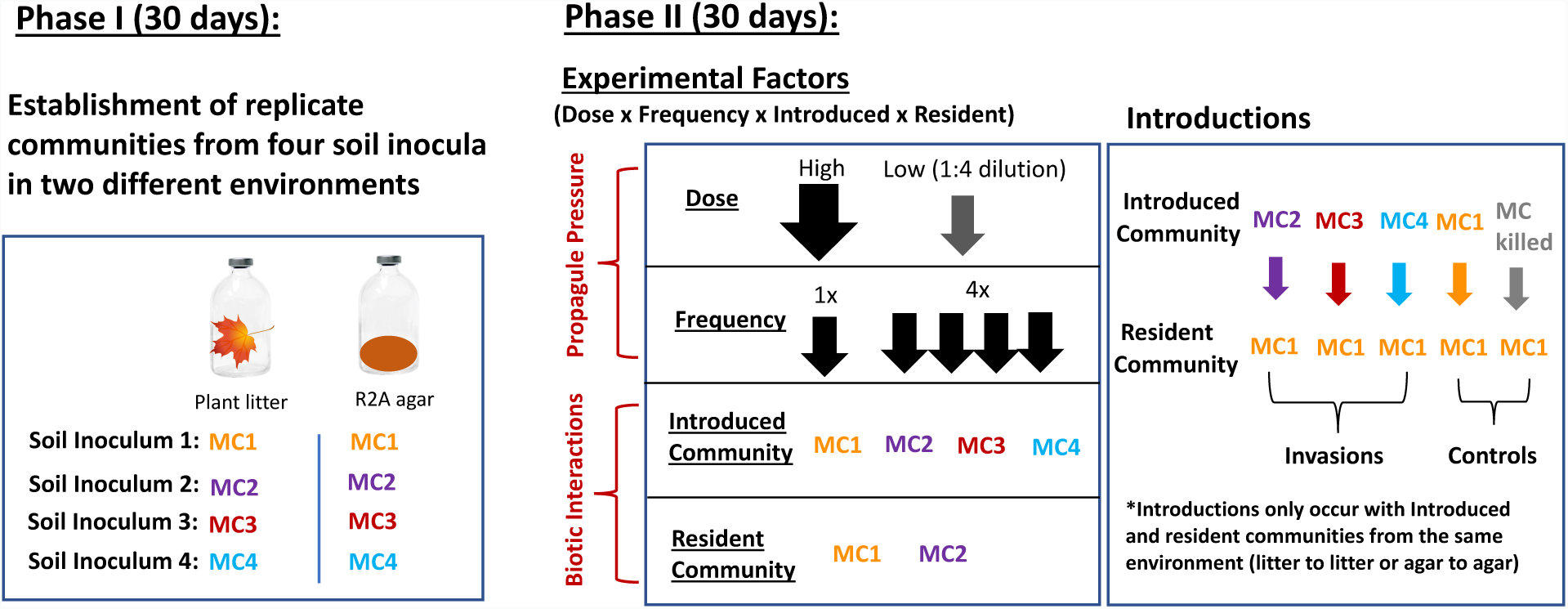
Experimental set-up used to test factors driving composition and functional outcomes of microbial community invasions. In *Phase I* four soil inocula were used to inoculate microcosms in order to establish many replicates of complex communities in plant litter and R2A agar substrates. In Phase II we conducted microbial community invasions, while varying four factors including dose, frequency, introduced, and resident communities (See methods section for details).

## Results

### Impacts of propagule pressure and biotic interactions in shaping microbial communities

With the goal of identifying common rules governing impacts of microbial invasions on community composition across ecosystems, we assessed the relative roles of propagule pressure and biotic interactions in driving microbial composition separately across the R2A agar and plant litter environments. Specifically, the impacts of invader community (MC1, MC2, MC3, MC4), resident community (MC1, MC2), dose (high, low), and frequency (high, low) were tested (Figure 1). In both environments, community composition was shaped by a combination of the treatment factors, where biotic interactions (invader and resident community) played a larger role than propagule pressure (dose and frequency) (Figure 2, Table S1). Biotic interactions also played a greater role than propagule pressure in driving bacterial and fungal richness (Figure 2, Table S1).

**Figure 2.**
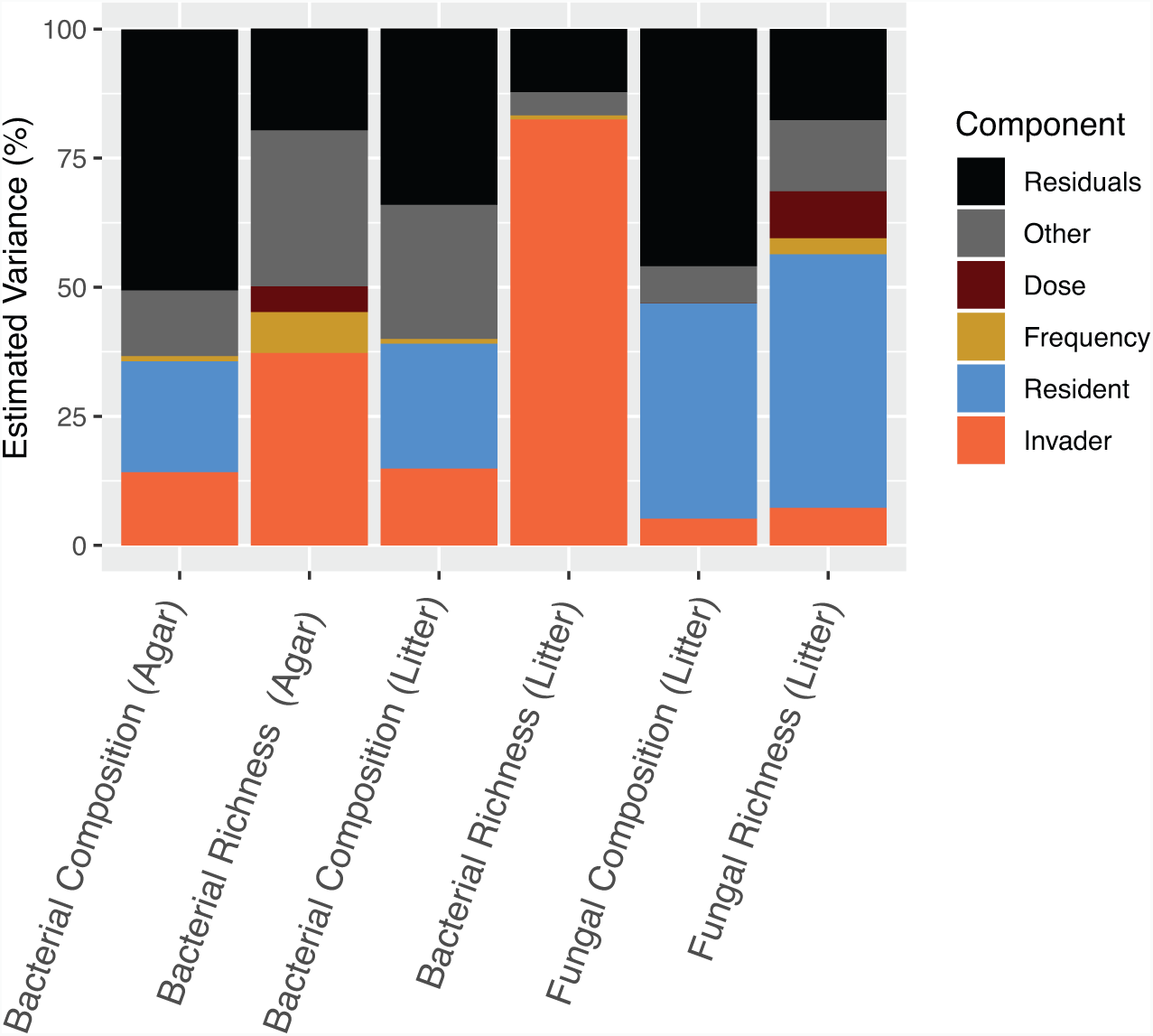
Impact of propagule pressure (dose and frequency) and biotic factors (resident and invader composition) in driving bacterial and fungal composition and richness in litter and agar environments. Estimated variance was computed on reduced ANOVA and PERMANOVA models. Only significant main factors are shown and ‘Other’ component is a sum of significant interaction terms. Complete statistics are in Table S1.

Within biotic interactions, the impact of invader versus resident community composition varied depending on the organism type. For bacteria invaders and residents both played a role in driving variation in community composition. 14% and 15% of estimated variation in composition was driven by invaders in the agar and litter respectively, while resident driven variation accounted for 22% in agar and 24% in litter (Figure 2, Table S1). In contrast, resident communities played a larger role than introduced communities in driving community composition for fungi in litter, contributing to 42% and 5% of estimated variance in composition respectively (Figure 2, Table S1). In the agar, fungal communities collapsed to below sequence detection limits. Both bacterial and fungal community composition were also impacted by interaction terms including a resident-by-invader community interaction (Table S1). Invaded bacterial community composition was significantly different from nutrient addition control (e.g. killed MC1-MC1) and resident-resident control (e.g. MC1-MC1) (pairwise permutation MANOVAs, Figure S1). We observed similar trends for richness, with a greater impact of the invaders on bacteria, and a greater impact of residents on fungi (Figure 2). For bacteria invader richness was a driving factor, while fungal richness was more linked to resident richness (Figure S2, Figure S3). Generally invaded communities had higher bacterial richness and diversity than nutrient addition control (e.g. killed MC1-MC1) and resident control (e.g. MC1-MC1) (Figure S3abde). Fungal richness and diversity were similar across invaded and control communities within a single resident community type (Figure S3cf).

Compared to biotic interactions, propagule pressure played a minor role in shaping the microbial communities, with larger impacts on richness than composition (Figure 2). For richness, the influence of invasion frequency and dose varied depending on organism and environmental type. For bacteria, in the agar both frequency and dose, as well as a frequency-by-dose interactions played a role in driving richness (Figure 2). Higher bacterial richness occurred in higher dose and higher frequency invasions in agar, but not in litter (Figure S4abde). For fungi, higher dose invasions led to higher richness, however, higher frequency invasions led to decreased richness (Figure S4cf).

As expected, overall the environment was a strong driver of composition. Bacterial communities differed by environment type (Figure S5a), but after 60 days bacterial richness was similar across the two environments (Figure S5b). Fungal communities collapsed in the agar, thus analyses for fungal communities was performed only in the litter environment.

### Quantifying invasion success and identifying invader taxa

To quantify the influence of biotic interactions in driving the outcome of invasion events, we identified the taxa that were invaders, non-invaders, resilient, non-resilient, or common to microbial communities. To do this we assessed Operational Taxonomic Unit (OTU) presence/absence across the 1) Invader Inoculum 2) Resident Control (resident-resident, killed-resident, and initial *Phase II* resident samples), and 3) Final Invaded communities (Figure 3, Table S2). On average bacterial communities contained 357 ± 67 OTUs (Table S2). Across the twelve invasion events, the distribution of OTUs across categories was highly consistent. For bacteria invaders comprised 9 ± 6% of the OTUs. This is likely a conservative estimate for overall invader numbers given that 20± 5% of bacterial OTUs were of undetermined origin (i.e. OTUs only found in the Final Invaded communities, which must have been undetected in either the Resident Control or Invader Inoculum communities) (Figure 3a,b). It is more likely they were undetected in the Invader Inoculum for the following reasons: 1) we sequenced more Resident Control as compared to Invader Inoculum samples (Table S2) as we wanted to limit the detection of false positives for invader taxa and; 2) the percentage of OTUs unique to Final Invaded communities was positively correlated (R^2^=0.6, p=0.04) with the average OTU richness of the Invader Inoculum (Figure S6a). Resilient bacteria comprised 25 ± 7% of bacterial OTUs (Figure 3). The percentage of resilient OTUs was negatively correlated with the average OTU richness of the Invader Inoculum (R^2^=-0.71, p=0.009) (Figure S6b). 14 ± 5% of bacterial OTUs were common (Figure 3, Table S1). Common bacterial OTUs increased as inoculum richness increased (R^2^=0.61, p=0.04, Figure S6c). On average fungal litter communities were comprised of 164±67 OTUs. The distribution of OTUs across categories was very similar to bacteria, with the exception of an increase in non-invasive fungi, which made up 19±6% of OTUs in fungi versus 7±3% of OTUs in bacteria (Figure 3).

**Figure 3.**
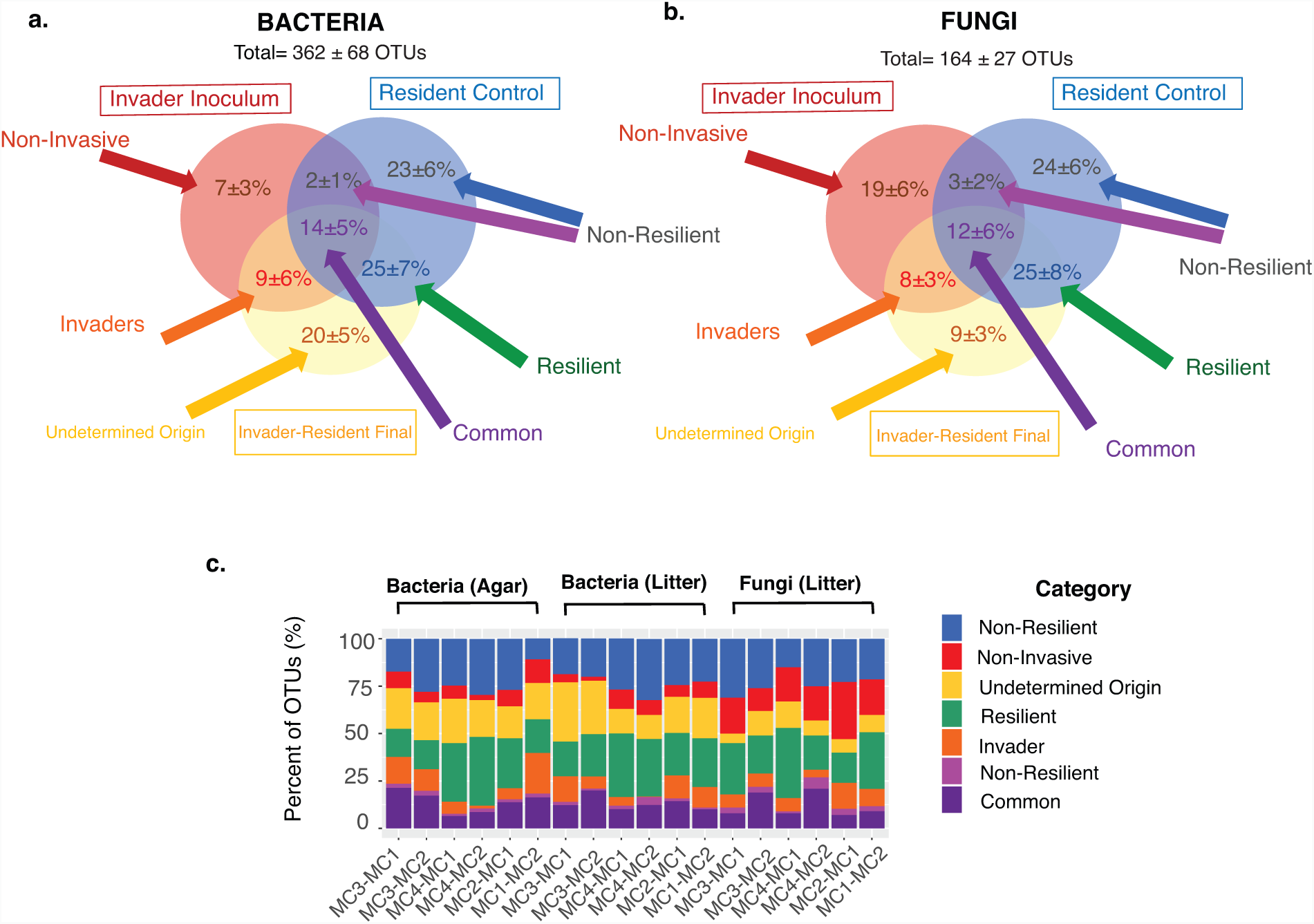
Percentage of OTUs that were invaders, resilient, common, non-invasive or non-resilient across the twelve invasion events. Average OTU distribution for a) Bacteria and b) Fungi. c) Distribution of OTUs across categories for each individual invasion event. (Data shown in Table S2).

For bacteria, 148 successful invaders were identified, and 96 were invaders in at least two invasion events (Dataset S1). The genera *Arthrobacter* (Actinobacteria) and *Microvirga* (Alphaproteobacteria) were invaders in the most events (4 out of 12). Overall 23 invaders were found in high abundance (High abundance=average OTU count >10 OTUs, Low abundance=average OTU count < 10 OTUs) in the final communities, where most of these (18 out of 23) started out in low abundance in invader inoculum communities. The invaders with the highest final abundance were primarily from the Bacteriodetes (Sphingobacteriia (n=13)) and Proteobacteria (Alphaproteobacteria (n=3) and Gammaproteobacteria (n=4)).

For fungi, 69 successful invaders were identified, and nine of these were invaders in at least two events (Dataset S2). These common invaders included two *Chaetomidium* genera, as well as *Christiansenia*, *Phaesophaeria*, *Nectaria, Peyronellaea*, *Giberella*, *Necocosmospora*, and *Rhodotorula* genera. Invaders were found in both high and low abundance in the invader inoculum, and almost exclusively were low abundance in the final communities. Only one fungal OTU (*Chalara* genera) was found in low abundance in inoculum (MC1), but high abundance in final communities (MC1-MC2).

### Impacts of propagule pressure and biotic interactions in driving variation in ecosystem functioning

Overall, total CO_2_ production across all invaded microcosms varied by 3.1-fold in the agar and 2.0-fold in the litter (Figure 4a), while DOC accumulation varied by 3.1-fold in the litter and 3.3-fold in the agar (Figure 4b). As with composition, functioning was primarily driven by biotic interactions (Figure 4c). Within biotic interactions, the impact of invader versus resident community on ecosystem functioning varied by environment (Figure 4c). In the agar the invader community accounted for 80% of variation in CO_2_ production and 40% of variation in DOC accumulation. By contrast, both the resident and invader communities in the litter were important to CO_2_ production, accounting for 27% and 17 % of estimated variation respectively. In the agar, some invaded communities produced significantly more CO_2_ compared to controls, while this was not the case in litter (Figure S7). However, in the litter environment, CO_2_ production did not vary across invaded and control communities (Figure S7). Overall, invader MC3 had the greatest impact on ecosystem functioning (Figure S7). In sum, these results suggest that the potential to use invasions to alter functioning may depend on both the environment and invader community.

**Figure 4.**
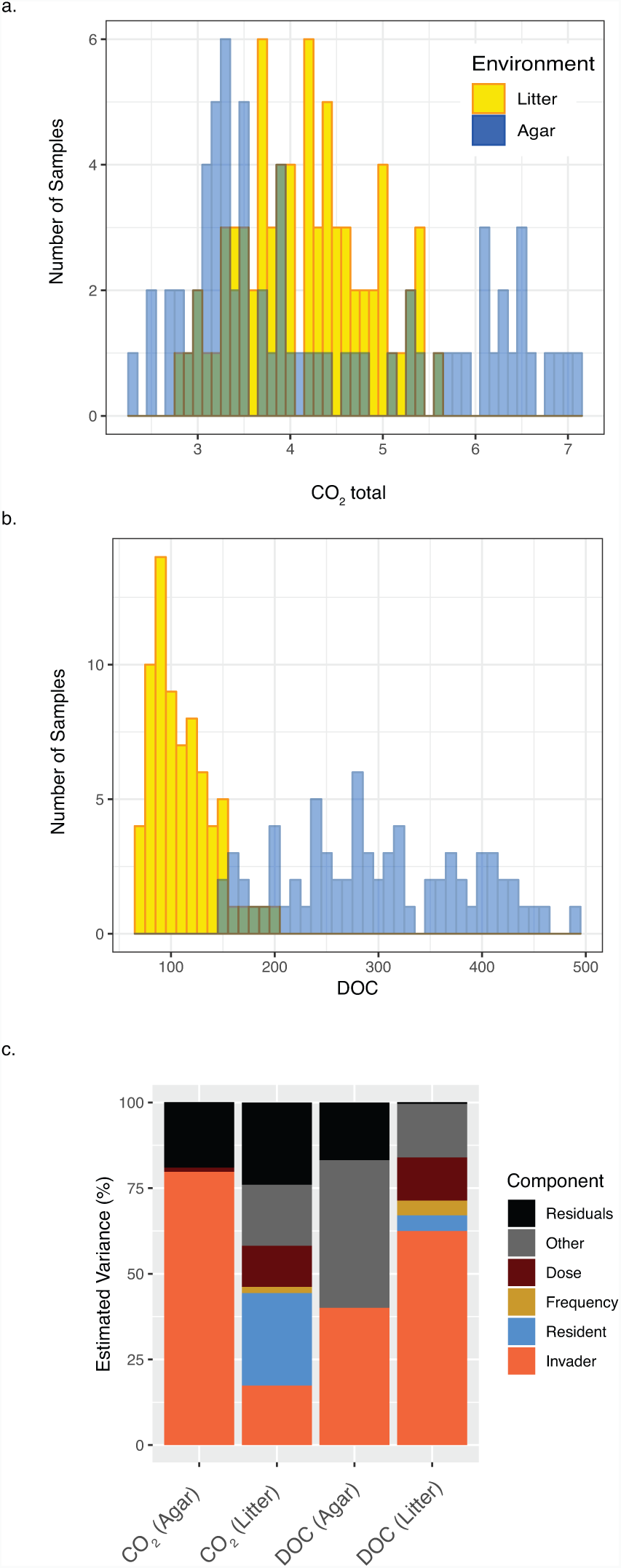
Histogram of the distribution of ecosystem functioning metrics, (a) CO_2_ production and (b) DOC accumulation, across *Phase II* invaded microcosms (n=144). Litter environment measures are shown in gold and media environment measures are shown in blue. (c) Impact of propagule pressure (dose and frequency) and biotic interactions (resident and introduced composition) in driving ecosystem functioning measured as total CO_2_ production and DOC accumulation. Estimated variance was computed on reduced ANOVA models. Only significant main factors are shown and ‘Other’ component is a sum of significant interaction terms. Complete statistics are in Table S3.

Propagule pressure did not impact functional outcomes in the agar, with the exception of the minor role of dose (1.2%) in driving variation in CO_2_ production. However, in the litter, dose played a role in driving variance in both CO_2_ production (12%) and DOC accumulation (12%) (Figure S2, Table S3), with increases in CO_2_ production and DOC accumulation in the high compared to low dose samples (Figure S8). Furthermore, frequency played a minor role in CO2 production (1.8%) and DOC accumulation (4.3%) in the litter, with four introductions having higher DOC accumulation than one introduction.

Across all samples final bacterial community composition was linked to both CO_2_ and DOC (PERMANOVA; Agar CO_2_: F_2,4_=1.9, p=0.004; DOC: F_2,4_=8.8, p=0.001; CO_2_-by-DOC: F_4,92_=1.3, p=0.028, Litter CO_2_: F_2,4_=3.3, p=0.001; DOC: F_2,4_=6.4, p=0.001; CO_2_-by-DOC: F_4,101_=1.4, p=0.062) (Figure 5a). Bacterial richness was positively correlated with cumulative CO_2_ production in both environments, although the correlation was stronger for agar than litter (R^2^=0.72, p<0.001 and R^2^=0.29, p=0.002, Figure S9a). Bacterial richness was negatively correlated with DOC in litter (R^2^=-0.63, p<0.001) and agar (R^2^=-0.4, p<0.001) environments (Figure S9c). Fungal community composition on litter was linked to CO_2_ production, but not DOC (PERMANOVA; CO_2_: F_2,4_=3.1, p=0.002; DOC: F_2,4_=1.7, p=0.1; CO_2_-by-DOC: F_4,71_=0.9, p=0.636) (Figure 5b). The differences in CO_2_ production across the fungal communities largely corresponded to the resident community composition. Fungal richness was negatively correlated with CO_2_ production (R^2^= −0.28, p=0.007, Figure S9b) but was not significantly correlated with DOC (Figure S9d).

**Figure 5.**
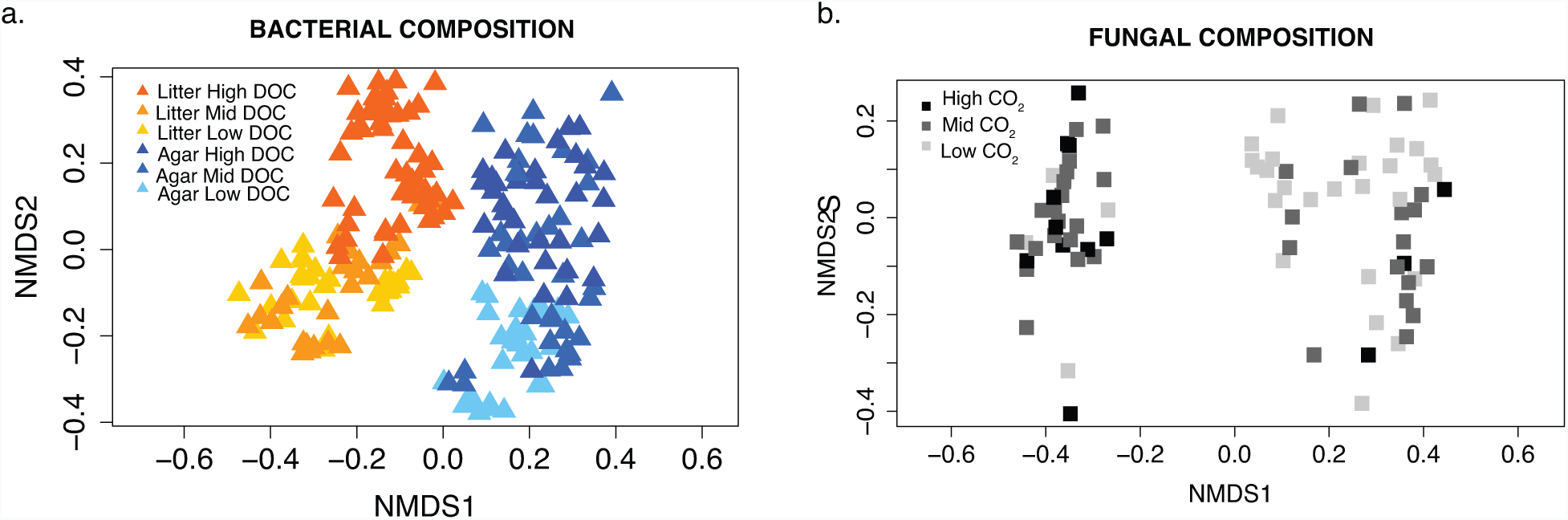
Nonmetric multidimensional scaling (NMDS) ordinations showing variability in (a) 16S bacterial community composition (16S) of final communities (n=212) and b) fungal community composition (LSU) of final communities (n=96) using the Bray-Curtis dissimilarity metric. Points are colored by the most significant correlation with functioning, for bacteria this is by high, mid and low DOC in each environment type, while for fungi by high, mid, and low CO_2_.

## Discussion

Discovering basic principles for successful microbiome engineering across systems is needed to increase the effectiveness of probiotics. To our knowledge this is the first study to test the relative importance of propagule pressure and biotic interactions in driving variation in microbial community composition and ecosystem functioning following a microbial community invasion event. In contrast to our hypothesis propagule pressure was less important than biotic interactions in driving variation in microbial composition and ecosystem functioning (Figure 2). In both macro- and micro-organism studies increasing the number of invading individuals and the number of repeated introductions leads to higher invasion success in both field and laboratory experiments (10, 15, 20, 21, 39, 48, 49). Invasions with large numbers of diverse taxa compared to single taxa may increase the importance of biotic interactions relative to propagule pressure. In this scenario residents must resist larger numbers of invaders and invaders must compete both with other invader taxa, as well as the established resident taxa. A few studies have found that groups of microbes have the potential to be both more robust and more productive than monocultures (50–52). With a diverse invader community, the probability of an introduced taxon occurring that is highly competitive and may establish regardless of propagule pressure increases. This is supported by the observation that invaders occurred in both high and low abundance in the invader inoculum (Dataset S1, Dataset S2). Overall, our results suggest that when engineering microbial communities with a goal of altering microbial composition, resources would be better spent focusing on characteristics of the microbes being introduced and of the microbes already present, rather than delivery dose and frequency.

Given the increasing evidence of strong links between microbial community composition and functioning (53–55), it is perhaps unsurprising that functioning was most strongly impacted by the same dominant factor --biotic interactions-- as community composition. Furthermore, the environment played an important role in shaping the impact of invaders on ecosystem functioning (Figure 4c). In the agar, the larger role of invader communities in driving changes in ecosystem functioning may be due to the loss of fungi, leading to a more prominent role of bacterial invaders. By contrast on the litter, fungi were likely the most important players in driving CO_2_ dynamics. Fungal composition on litter was most significantly associated with respiration (Figure 5b) and the resident litter communities played the greatest role in determining both fungal composition and CO_2_ production (Figure 2, Figure 4c). Higher growth rate and turnover on agar compared to litter may also have contributed to larger invader impacts on ecosystem functioning, as bacterial invaders were equally important in driving composition in agar and litter, but had a much greater impact on ecosystem functioning in agar. Overall, our results suggest that potential exists for community invasions to alter ecosystem functioning, but the environment matters likely through its role in changing microorganism interactions.

While biotic interactions were key, the relative impact of invader compared to resident communities varied by organism type (Figure 2). Previous research has largely focused on the role of increased resident community richness and diversity on reducing invasion success (1, 11, 24–27). In part this is because current engineering applications routinely introduce individual or a few microbial taxa (11, 56–58). With community invasions, invaders were more important in driving bacterial communities, whereas residents played a larger role in shaping fungal communities (Figure 2). In addition, bacteria were better invaders than fungi (Figure 3). Other studies looking at the effects of abiotic (environmental) perturbations on microbial communities have found that bacterial communities are generally less resistant to change than fungal communities (55, 59). More generally, bacterial and fungal community assembly has been found to differ (60–62). Factors such as growth rate, growth habit (unicellular vs filamentous), and resource utilization breadth have also been hypothesized to play a role in differences in bacterial versus fungal establishment (55, 63). We expect these factors are likely important in our system as well. Our results suggest that engineering fungal communities may be more difficult than bacterial communities.

An advantage of conducting introductions with complex microbial communities is that it allows identification of a suite of successful invaders in a single experiment, which facilitates the search for common invader characteristics. The ability of microbes to establish in environments with resident communities is key to success in engineering applications. However, little is known about invasive microbes or their traits (64). Most previous experimental microbial invasion studies have picked a single to a few invaders of interest *apriori* (20). Characteristics, such as dispersal ability, reproductive strategy, and growth form have been linked to invader success in macro-organisms (65–67). We identified a number of successful invader microbes. In particular Sphingobacteria (Bacteriodetes) stood out as an invader by contributing approximately half of the invader taxa found in the highest abundance in final communities. Sphingobacteria are environmental bacteria capable of producing sphingolipids, a relatively uncommon microbial trait (68). Sphingolipids have been shown to play an important role in promoting bacterial virulence and enhancing survival during stress (68, 69). At a broad level, invasive fungal taxa identified in this study have been linked to pathogens. Fungal invaders predominantly included genera known to contain members that are pathogens, including plant pathogens (Nectria, Giberella, Necocosmospora, Peyronellaea, Chalara, Phaesophaeria), a fungal parasite (Christiansenia), and a genus known to contain human pathogens (Rhodotorula). Common pathogen traits such as ability to adhere to host cells, motility, toxin production, and/or antibiotic resistance (70) may be linked to their invasiveness. Future research is needed to assess trade-offs in traits linked to invasion success.

We acknowledge a number of limitations to this study. Using soil suspensions to create complex microbial communities leads to a loss of information of all the individual taxa in the system. This approach thus lacks control of factors such as microbial richness or diversity that may alter invasion dynamics (17, 19, 71). In addition, impacts of invasions were measured at a single timepoint, but impacts may change over time. For example, invaders might persist below detection limits until an opportunity arises in the process of community succession and environmental modifications for them to flourish. Also, to consider is that identified invaders were found primarily in low abundance and thus over a longer timeframe they could potentially be lost. Overall, this points to the need for experiments tracing community succession after invasion events. Tracing succession would not only provide greater insight into biotic filtering, but may also reveal inflection points in the relative importance of biotic filtering and propagule pressure if these factors exert impacts over different timescales.

The goal in engineering microbial communities is to alter microbiome functioning via introduction of invaders. Our results suggest that a decision tree for probiotic design should start by considering characteristics of the target environment that may influence organism interactions. The next step is to consider biotic interactions, in particular the ability of an invader community to establish which varies by organism type (i.e. bacteria versus fungi).

Delivery parameters (dose and frequency) may be considered as the last measure to increase invasion success. Probiotic development is a booming industry, where markets values for human and animal products alone are expected to exceed $50 billion by 2022 (31), while interest in plant probiotics is also increasing (32–34). A better understanding of fundamental principles that enhance the establishment and resilience of microbial inoculants has the potential to increase successful engineering of microbial communities for applications in human health, agriculture, bioenergy, and biotechnology.

## Methods

### Experimental Approach

We used a laboratory microcosm experiment, where we manipulated four factors: inoculation dose, inoculation frequency, introduced microbial community and resident microbial community across two environments. During *Phase I* of the experiment we established diverse model microbial communities in microcosms, while in *Phase II*, we conducted experimental manipulations adding introduced microbial communities at different doses and frequencies to established resident microbial communities (Figure 1).

#### Phase I

We created two environmental types in microcosms, a relatively nutrient rich environment (R2A agar medium containing diverse carbon sources) and a relatively nutrient poor environment (sterilized ground pine (*Pinus Ponderosa*) litter on sterile sand). We then created model microbial communities by inoculating the microcosms with suspensions of soil microbes from four soil samples from disparate locations, (35°01’49.4"N 106°03’03.9"W, 38°18’36.0"N 109°16’48.0"W, 35°58’41.430"N 79°05’39.087"W, 35°06’14.4"N 106°36’17.2"W), hereafter referred to as model communities MC1, MC2, MC3, and MC4. A microbial slurry was created by diluting soil in a PBS and NH_4_NO_3_ (1 mg/mL) solution. Two initial soils were randomly chosen to create resident microbial community types (MC1 and MC2). 1.3 mL of the microbial slurry of each resident microbial community type was used to inoculate 60 microcosms of each environmental type, for a total of 240 microcosms. Microcosms were covered with aluminum foil and placed in a 25°C incubator for one month. This time period was chosen to allow the microbial communities from soil to establish in the novel environment. 0.5 mL of sterilized H_2_O was added to each microcosm weekly to prevent microcosms from drying out. At the end of one month, three microcosms of each type (environment-by-resident community) were destructively sampled for DNA sequencing. To do so, 5 mL of H_2_O was added to the microcosms. For R2A microcosms a scraper was used to scrape the biofilm into solution. Microcosm material was gently vortexed for 5 seconds and swirled for 30 seconds to homogenize the mixture. We then sampled 2 mL, which was saved at −80 °C until subsequent DNA extractions and sequencing.

All four model communities, MC1, MC2, MC3, and MC4, were used to create introduced microbial communities. At the start of *Phase I* for each of the four communities, we used 1.3 mL of each slurry to inoculate 15 microcosms of each environmental type. These microcosms received the same treatment as resident community microcosms during *Phase I*. However, at the end of *Phase I*, these microcosms were all destructively sampled to create introduced community inoculum. Samples were homogenized, using the same method described above for the resident community DNA sampling. However, here the one-month slurry from the 15 microcosms of each microbial community-by-environment type was combined to create 8 inoculation slurries. During this combination step in the pine litter microcosms we filtered out the pine litter using a 40 µM filter. For R2A agar introduced communities, microbial abundance was estimated using optical density (OD) measurements and the communities were diluted to a common OD measurement.

To create different dose treatments, for each microbial community-by-environment type (n=8) we split the sample into three portions. One aliquot of the one-month slurry (standardized for R2A media) was kept for the high dose treatment. The second aliquot was diluted to a 1:4 ratio and used as the low dose treatment. This dilution ratio was chosen to minimize the impacts of dilution on composition in the complex communities and to parallel the frequency treatments (1x, 4x). The third aliquot was autoclaved in order to use killed microbial communities as a control treatment. This control dose treatment was used to make sure that effects of invasions were not just due to additional nutrient inputs. Subsamples of the *initial Phase II* introduced microbial communities were collected and stored, along with the *initial Phase II* resident microbial communities at −80 °C until subsequent DNA extractions and sequencing.

#### Phase II

We used a crossed experimental design to test the effects of introduction frequency (1x or 4x) and introduction dose (high or low (1:4) or killed) in determining microbial community composition and functioning (see Figure 1). Microbial community introductions were performed by adding 0.5 mL of an introduced model community (MC1, MC2, MC3, MC4) into each of the resident community microcosms (MC1, MC2), with three replicates for each treatment type (n=240 total microcosms). Microcosms with established resident communities received a high dose, low dose, or killed dose of introduced communities on day 1 of *Phase II*. Microcosms were then sealed and placed in a 25 °C incubator. Approximately weekly, on day 9, day 16, and day 23, the 4x frequency treatment microcosms received 0.5 mL of the same introduced model community as on day 1, while the 1x frequency treatments received 0.5 mL of H_2_O for the pine litter environment or R2A media for the R2A environment. For the killed-dose treatment, introductions were performed only at the 4x frequency, as the goal was to use this as a control. Introduced communities were stored at 4 °C in-between introductions in order to limit changes in communities, however we acknowledge that communities at 4°C may not be static during a 23-day incubation. Once sealed, liquid introductions to microcosms were delivered via a sterilized syringe. CO_2_ was measured using an Agilent Technologies 490 Micro Gas Chromatographer (GC) on days 2, 5, 9, 16, 21, and 30. Headspace was replaced with ambient sterile-filtered air after measurements to prevent oxygen depletion and CO2 build-up. At 30 days after the final CO_2_ measurement, microcosms were destructively sampled using the same approach as in *Phase I*. 1.5 mL of liquid from each microcosm were archived at −80°C for DNA extraction. The remaining 3.5 mL was filter-sterilized (0.2 uM filter; item detail) and stored at - 20°C for dissolved organic carbon (DOC) measurements. DOC concentration was measured on an OI Analytical model 1010 wet oxidation TOC analyzer (Xylem Inc., Rye Brook, NJ, USA).

### DNA extractions and microbial community sequencing

DNA extractions on samples from introduced communities and resident communities at start of *Phase II*), as well as all microcosm communities at the end of *Phase II* were performed with a PowerSoil DNA extraction kit (Mobio). The standard protocol was used with the exception that 0.5 mL of the homogenized liquid sample was used per extraction. Taxonomic profiling was performed by sequencing bacterial 16S rRNA and fungal 28S rRNA genes. The V3-V4 region of bacterial (and archaeal) 16S rRNA genes was amplified using primers 515f-R806 (72) and the D2 hypervariable region of fungal 28S rRNA gene was amplified using primers LR22R (73) and LR3 (74). PCR amplifications for bacteria and fungi were performed using a two-step approach (73). In the first PCR, sample barcoding was performed with forward and reverse primers each containing a 6-bp barcode; 22 cycles with an annealing temperature of 60°C were performed (75). The second PCR added Illumina adaptors over 10 cycles with an annealing temperature of 65°C. Amplicon clean-up was performed with a Mobio UltraClean PCR clean-up kit, following manufacturer’s instructions with the following modifications: binding buffer amount was reduced from 5X to 3X sample volume, and final elutions were performed with 50 µl Elution Buffer. Following clean-up, samples were quantified with an Invitrogen Quant-iT^TM^ ds DNA Assay Kit on a BioTek Synergy HI Hybrid Reader. and pooled at a concentration of 10 ng per sample. A final clean-up step was performed on pooled samples using the Mobio UltraClean PCR clean-up kit. Samples were sequenced on an Illumina MiSeq platform with PE250 chemistry at Los Alamos National Laboratory. Unprocessed sequences are available through NCBI’s Sequence Read Archive (PRJNA557183).

### Microbial community sequence analysis

Bacterial and fungal sequences were merged with PEAR v 9.6 (76), quality filtered to remove sequences with 1% or more low-quality (q20) bases, and demultiplexed using mothur (77) allowing no mismatches to the barcode or primer sequence. Further processing was undertaken with UPARSE (78). Sequences with an error rate greater than 0.5 were removed, remaining sequences were dereplicated, singletons were excluded from clustering, OTU clustering was performed at 97% and putative chimeras were identified *de novo* using UCHIME (78). Previous analyses have shown congruent ecological patterns with use of OTUs vs ESV’s for delineating microbial taxa (79). Furthermore, OTU clustering at 97% provides a more conservative estimate of overlaps between introduced and resident taxa. Bacterial and fungal OTUs were classified using the Ribosomal Database Project (RDP) classifier (80). Using the OTU matrices for final communities (n=240), we generated rarefied composition distance matrices for bacteria and fungi randomly drawing the lowest common number of sequences from each sample, n= 1020 and n=1262, respectively. We then calculated a Bray-Curtis distance matrix for bacteria and for fungi (81). In addition, richness and Shannon diversity of rarified OTU matrices was also calculated (82). Trends were similar for richness and diversity across all analyses.

In addition to analyses at the community level, we also looked at how introductions altered microbial composition at the taxa (OTU) level. Here we started with an OTU table including both initial *Phase II* and final *Phase II* communities. The four initial Phase II inoculum communities for both bacteria and fungi in the two different environment types were significantly different from each other (pairwise permutation MANOVAs). Community composition of inoculum is shown in (Figure S10). Replicate resident communities within an environment from the initial *Phase II* sampling were more similar to each other than to other resident communities in that environment (Figure S11). The four complex model microbial communities contained both common and unique taxa, therefore we looked at the distribution of taxa across the different initial communities, and how that distribution changed with the addition of each invader community. Here, we split our analyses by environmental type, as our community analyses showed that the environment played a large role in structuring final microbial composition. For each invader-by-resident community combination, we compared the presence/absence of OTUs in the final invader-resident samples (Invader-Resident), to OTUs in the introduced inoculum samples (Invader Initial) and the resident control samples (Resident Control). The Resident Control included resident-resident final samples, resident-killed final samples, and initial resident samples. Overall, this led to analysis of 12 unique introduction events, including 6 per environment type (MC1 into MC2, MC3 into MC2, MC4 into MC2, MC2 into MC1, MC3 into MC1, MC4 into MC1). For each event, we calculated the percentage of the total OTUs found in each category (Resident Control Only, Invader Inoculum Only, Invader-Resident Final Only, Resident Control + Invader-Resident Final, Invader Inoculum + Invader-Resident Final, Invader Inoculum + Resident Control, All).

### Statistical analysis

First, using the end communities for the Phase *II* experimental treatments, we tested the impact of invaders on community composition and ecosystem functioning by comparing the influence of invader (e.g. MC2-MC1) compared to control treatments (e.g. Killed MC1-MC1 and MC1-MC1) on community composition and ecosystem functioning across two environments (agar and litter). Here we used a one-way ANOVA for univariate metrics (i.e. CO_2_ production, DOC production, richness, Shannon diversity) and pairwise permutational MANOVAs for multivariate metrics (i.e. bacterial and fungal composition).

Next, using only final invaded end communities (n=144) excluding controls, see Figure 1), we looked at what factors most impacted microbial composition and ecosystem functioning following invasion events. Here, our *Phase II* experimental treatments included four factors invader community (MC1, MC2, MC3, MC4), resident community (MC1, MC2), dose (high(no dilution), low (1:4 dilution)), and frequency (1x addition, 4x addition), which we examined across two environment types (agar, litter). To test the impact of treatment factors and estimate the variance explained by each treatment in driving variation in univariate metrics (i.e. CO_2_ production, DOC production, richness, Shannon diversity) we used a multi-factorial ANOVA design with all four manipulated variables as main fixed factors. We tested the effects of the main factors as well as interaction terms (Dose * Frequency * Invader * Resident). Analyses were performed on full models and then reduced models were run with only the significant factors. Analyses were performed separately for each environmental type. The ANOVA analyses were conducted in the R software environment (83). To assess the contribution of treatments in driving variation in bacterial and fungal community composition, we performed a permutational multivariate analysis of variance (PERMANOVA)(84), using the same factors as with the univariate tests. Using results from the reduced models, we estimated the percent of variation that could be attributed to each significant term for both the ANOVA (85) and PERMANOVA (as described in (86)) analyses. Using community composition data from all samples including controls (n=240), we tested the correspondence between microbial (fungal and bacterial) community and functional metrics (CO_2_ and DOC) was performed using Pearson’s correlations for univariate metrics (richness, diversity) (Rcorr package). In addition, in order to link microbial community composition and functioning, we grouped samples into high (1/3), mid (1/3), and low (1/3) categories based on DOC and CO_2_ values. We ran a two-way PERMANOVA using DOC and CO_2_ categories as factors, including a DOC-by-CO_2_ interactions term.

## Acknowledgements

Thank you to Kendra Walters, Marie Kroeger, Joany Babilonia, Jeff Heikoop, and Brent Newman for comments of previous versions of this manuscript. Thank you to Andreas Runde for laboratory assistance. This work was supported by a Los Alamos National Laboratory Postdoc Fellowship to MBNA and by SFA grant 2019LANLF260 from the US Department of Energy Office of Biological and Environmental Research to JD.

## Author contributions

M.B.N.A and J.D. designed research. M.B.N.A. and L.V.G.G. performed the research. M.B.N.A. analyzed the data. M.B.N.A., J.D, and S.S. wrote the paper.

